# Single-cell transcriptomics predict novel potential regulators of acute epithelial restitution in the ischemia-injured intestine

**DOI:** 10.1101/2024.06.28.601271

**Authors:** Elizabeth C. Rose, Jeremy M. Simon, Ismael Gomez-Martinez, Scott T. Magness, Jack Odle, Anthony T. Blikslager, Amanda L. Ziegler

**Affiliations:** North Carolina State University, College of Veterinary Medicine, Department of Clinical Sciences, Raleigh, NC, USA; Department of Genetics, University of North Carolina at Chapel Hill, Chapel Hill, NC, USA; UNC Neuroscience Center, University of North Carolina at Chapel Hill, Chapel Hill, NC, USA; Bioinformatics and Analytics Research Collaborative, University of North Carolina at Chapel Hill, Chapel Hill, NC, USA; North Carolina State University, College of Agriculture and Life Sciences, Department of Animal Science, Raleigh, NC, USA

**Keywords:** intestinal ischemia, restitution, barrier function

## Abstract

Intestinal ischemic injury damages the epithelial barrier predisposes patients to life-threatening sepsis unless that barrier is rapidly restored. There is an age-dependency of intestinal recovery in that neonates are the most susceptible to succumb to disease of the intestinal barrier versus older patients. We have developed a pig model that demonstrates age-dependent failure of intestinal barrier restitution in neonatal pigs which can be rescued by the direct application of juvenile pig mucosal tissue, but the mechanisms of rescue remain undefined. We hypothesized that by identifying a subpopulation of restituting enterocytes by their expression of cell migration transcriptional pathways, we can then predict novel upstream regulators of age-dependent restitution response programs. Superficial mucosal epithelial cells from recovering ischemic jejunum of juvenile pigs were processed for single cell RNA sequencing analysis, and predicted upstream regulators were assessed in a porcine intestinal epithelial cell line (IPEC-J2) and banked tissues. A subcluster of absorptive enterocytes expressed several cell migration pathways key to restitution. Differentially expressed genes in this subcluster predicted their upstream regulation included colony stimulating factor-1 (CSF-1). We validated age-dependent induction of *CSF-1* by ischemia and documented that CSF-1 and CSF1R co-localized in ischemic juvenile, but not neonatal, wound-adjacent epithelial cells and in the restituted epithelium of juveniles and rescued (but not control) neonates. Further, the CSF1R inhibitor BLZ945 reduced restitution in scratch wounded IPEC-J2 cells. These studies validate an approach to inform potential novel therapeutic targets, such as CSF-1, to improve outcomes in neonates with intestinal injury in a unique pig model.

**NEW & NOTEWORTHY:** These studies validate an approach to identify and predict upstream regulation of restituting epithelium in a unique pig intestinal ischemic injury model. Identification of potential molecular mediators of restitution, such as CSF-1, will inform the development of targeted therapeutic interventions for medical management of patients with ischemia-mediated intestinal injury.

## INTRODUCTION

The small intestine is a crucial segment of the gastrointestinal tract in which a single layer of epithelial cells forms a selectively permeable barrier. This barrier permits absorption of ingested nutrients while maintaining a physical barricade from harmful, intralumenal stimuli such as bacteria and other microbes. When this epithelial barrier is compromised through epithelial cell loss, which may occur consequent to infection, mechanical damage, ischemic-injury and various other etiologies, those bacterial agents gain direct access to the host’s tissues and vasculature. Consequently, the host is at high risk for life-threatening sepsis unless that intestinal epithelial cell barrier is rapidly restored.

Early barrier restoration relies on epithelial restitution, which is the migration of viable, superficial epithelial cells across the denuded mucosal surface to reseal that physical barrier. As described in previous studies, restituting intestinal epithelial cells are characterized by a robust change in cellular morphology as they transition from their homeostatic, columnar shape to a flattened, almost squamous appearance.(1–3) To study restitution, our laboratory has developed a highly-translational pig model of ischemia-mediated loss of the intestinal epithelial barrier.(4, 5) Pigs share striking similarities in gastrointestinal anatomy, physiology, nutrition and microbiota with humans. Consequently, they are powerful models of myriad gastrointestinal diseases, including ischemic injury and restitution-mediated repair.(6–10) Our recent studies have identified an age-dependent difference in epithelial restitution responses in a highly relevant early-life pig model of intestinal ischemia and repair. We found that neonatal (nursing) pigs lack the restitution response while juvenile (weaned) pigs demonstrate robust restitution, and these slightly older pigs’ mucosal microenvironment can rescue the restitution phenotype in the neonatal pigs.(11, 12) Despite the critical role of these restituting cells in acute barrier restoration and therefore maintenance of intestinal health, much more is yet to be learned about the intercellular, molecular-level mechanisms that define and regulate this phenotypic change. Therefore, the overarching purpose of the current study was to identify the transcriptional profile that defines the restituting epithelial cell phenotype in the recovering jejunum of juvenile pigs and use this transcriptional profile to predict upstream regulators of the restitution phenotype. Identification of these transcriptomic signatures and its upstream regulators can inform targets for ongoing study with the goal of identifying novel regulators of restitution and thus potential novel therapeutic interventions to enhance intestinal epithelial barrier repair in vulnerable neonates.

A major limitation in further characterizing the restituting phenotype has been inadequate transcriptomic resolution of target cell populations. This limitation was grossly lessened, however, with the advent of single-cell RNA sequencing (scRNAseq). scRNAseq provides unprecedented transcriptomic profiling at cellular resolution, thereby providing the ability to further define cellular phenotypes and predict regulatory mechanisms that promote these phenotypes. Few studies have already defined methods through which resident cell populations in healthy gastrointestinal segments have been isolated and characterized.(13–15) Here, we successfully isolate and identify 8 distinct cell populations from the ischemia-injured jejunum of juvenile pigs. Based on gene expression characteristics of each cell type’s transcriptomic profile, we identified the restituting cell population, and highlight and preliminarily validate a potential upstream regulator of restitution, colony-stimulating factor-1 (CSF-1). Our results can inform ongoing investigations of potential novel therapeutic targets for patients with intestinal barrier injury.

## MATERIALS & METHODS

### Animals

All procedures were approved by the NC State University Institutional Animal Care and Use Committee. Yorkshire cross pigs 2- or 6-weeks-of-age of either sex were either euthanized at an academic commercial production facility for immediate tissue collection or were transported to a large animal surgical facility for surgical modeling experiments as described.

### Surgical Ischemic Injury Model

Pigs were sedated using xylazine (1.5 mg/kg) and ketamine (11 mg/kg). Anesthesia was induced with isoflurane vaporized in 100% oxygen via face mask, after which pigs were orotracheally intubated for continued delivery of isoflurane to maintain general anesthesia. Pigs were placed on a water-circulated heating pad and intravenous fluids were administered at a maintenance rate of 15 mL • kg^-1^ • h^-1^ throughout surgery. The distal jejunum was accessed via midline incision and 5- to 10-cm-long loops were isolated and subjected to 30 minutes of ischemia via ligation of local mesenteric vasculature with 2-0 braided silk suture. Within all experiments, ischemia-injured intestinal loops were resected and placed in oxygenated (95% O_2_/5% CO_2_) porcine ringer solution for transportation and subsequent *ex vivo* recovery. At the time of loop resection, pigs were humanely euthanized under general anesthesia with an overdose of pentobarbital. For *ex vivo* recovery, intestinal loops were opened and briefly rinsed in porcine Krebs-Ringer solution to remove intestinal contents from the mucosal surface. The rinsed intestinal tissue was then pinned to a dissection tray and bathed in warmed, oxygenated (95% O_2_/5% CO_2_) porcine ringer solution supplemented with 10 mM glucose for 30 minutes to permit epithelial recovery.

### Villus epithelial cell sloughing and dissociation

Following 30 minutes of *ex vivo* recovery, intestinal tissues were rinsed in Dulbecco’s phosphate-buffered saline (dPBS) at 4°C for 5 minutes to remove cations. Representative sections were retained for histology prior to cell sloughing. Rinsed tissues were incubated in ice-cold chelating buffer + 100 μmol/L Y-27632 for 20 minutes at 4°C.(16) Tissues were then transferred to fresh chelating buffer + 100 μmol/L Y-27632, incubated in a 37°C water bath and shaken to remove superficial villus epithelium. This experiment was repeated with 3 pigs, and high yield washes from all 3 animals were pooled into one tube. Representative sections of the remaining intestinal tissue were retained for histology. Villus epithelium was dissociated to single cells using 4 mg/mL Protease VIII in dPBS + Y-27632 on ice for approximately 30 minutes with trituration via a P1000 micropipette every 5 minutes. Cells were checked under a light microscope and then filtered.

### Sample preparation and single cell RNA sequencing

Single cells were washed with dPBS + Y-27632, resuspended in Advanced dPBS + 1% bovine serum albumin + Y-27632 for sorting on a Sony Cell Sorter SH800Z (Sony, Tokyo, Japan) to enrich for live single epithelial cells. Of the 8.3 million cells sorted, there were 147,000 cells that passed Annexin V live/dead and EpCAM parameters. These cells were loaded onto the Chromium Next GEM Single Cell 3’ GEM, Library and Gel Bead Kit v3.1 (PN-100012, 10x Genomics, Pleasanton, CA) for complementary DNA library preparation. Sequencing was performed on an Illumina NextSeq 500 (Illumina, San Diego, CA).

### Sequencing data analysis

Raw reads were processed and quantified using alevin v2.0.1 with Ensemble gene annotations for *S. scrofa* (v11.1.105), retaining all possible cell barcodes with “—keepCBFraction 1”.(17) Data were imported into R studio using tximport and converted into a Seurat object restricting to cells with at least 100 genes detected, and genes expressed in at least 100 cells.(18) Cells with high mitochondrial contribution were removed using MiQC, then an additional filter was applied to retain cells with at least 500 genes detected and 1000 UMIs, resulting in 1,359 cells and 13,337 genes.(19) Counts were then log-normalized and scaled, and dimensionality was reduced using the top 50 principal components. Cell types were discovered using Louvain-Jaccard clustering with multilevel refinement (algorithm=2, resolution=2.5). Clusters were visualized using UMAP and marker genes that characterized each cluster were detected using Seurat and Presto. Pathway enrichment analyses were performed using g:Profiler and (IPA^®^, QIAGEN Redwood City, www.qiagen.com/ingenuity). (20) Dendrograms were drawn based on 1-Pearson correlation distance of the top 50 most significant principal components. Dot plots were generated by running the DotPlot function in the Seurat (v4.3.1) R package.(21) Differential gene expression analysis was performed between cluster 0 and cluster 1. These data were then used as input for Qiagen Ingenuity Pathway Analysis (IPA^®^) software in order to assess enriched pathways, predicted upstream regulators and interaction networks. Gene expression in homogenates was presented using data which is publicly available on Gene Expression Omnibus (https://www.ncbi.nlm.nih.gov/geo/, accession #GSE212533) from prior studies.(12)

### Light microscopy

Tissues were fixed for 24 hours in 10% neutral-buffered formalin at room-temperature immediately following ischemic injury or after 120 minutes *ex vivo* recovery. Formalin-fixed tissues were transferred to 70% ethanol and then paraffin-embedded, sectioned (5 μm) and stained with hematoxylin and eosin for histologic analysis. Representative photomicrographs were captured with a Keyence BZ-X800E microscope.

### Immunofluorescence

Paraffin sections banked from prior rescue experiments reported by Ziegler et al. in PLoSOne in 2018 (11) were deparaffinized and washed 3 times in PBS to rehydrate the tissue. Tissues were cooled for 20-minutes at room temperature then placed in PBS-0.3% Triton −100 solution for 20-minutes to permeabilize the tissues. Tissues were washed twice in PBS and incubated in blocking solution (Dako, Carpinteria, CA, USA) for 1 hour at room temperature. Slides were incubated in the dark overnight at 4°C in rabbit anti-M-CSF polyclonal antibody (ThermoFisher PA542558, RRID:AB_260957) or mouse anti-CD115 monoclonal antibody (ThermoFisher MA528453, RRID:AB_2745415) at a dilution of 1:00 in diluent (Agilent Cat# S0809). After washing slides 3 times in PBS, slides were incubated for 1 hour at room temperature in the dark in donkey anti-rabbit conjugated to Alexa Fluor 594 (Thermo Fisher A21207; RRID:AB_141637) or goat anti-mouse conjugated to Alexa Fluor 488 (Thermo Fisher A11029; RRID:AB_2534088) diluted 1:500 in diluent (Agilent Cat# S0809). Tissues were counterstained with the nuclear marker 4′,6-diamidino-2-phenylindole (DAPI, Invitrogen Cat# D1306) at a dilution of 1:1000 in diluent (Agilent Cat# S0809) at room temperature. Slides were washed three times in PBS then mounted under a coverslip in aqueous mounting medium (Agilent Cat# S3025). Images were captured using a Keyence BZ-X800 All-in-One microscope (Osaka, Japan) using a 20x objective lens and images were processed using Keyence Software BZ Analyzer. One tissue per treatment was incubated in secondary antibody mixture with no primary antibody added serving as negative controls to ensure antibody specificity (Supp Fig 1).

**Figure 1.**
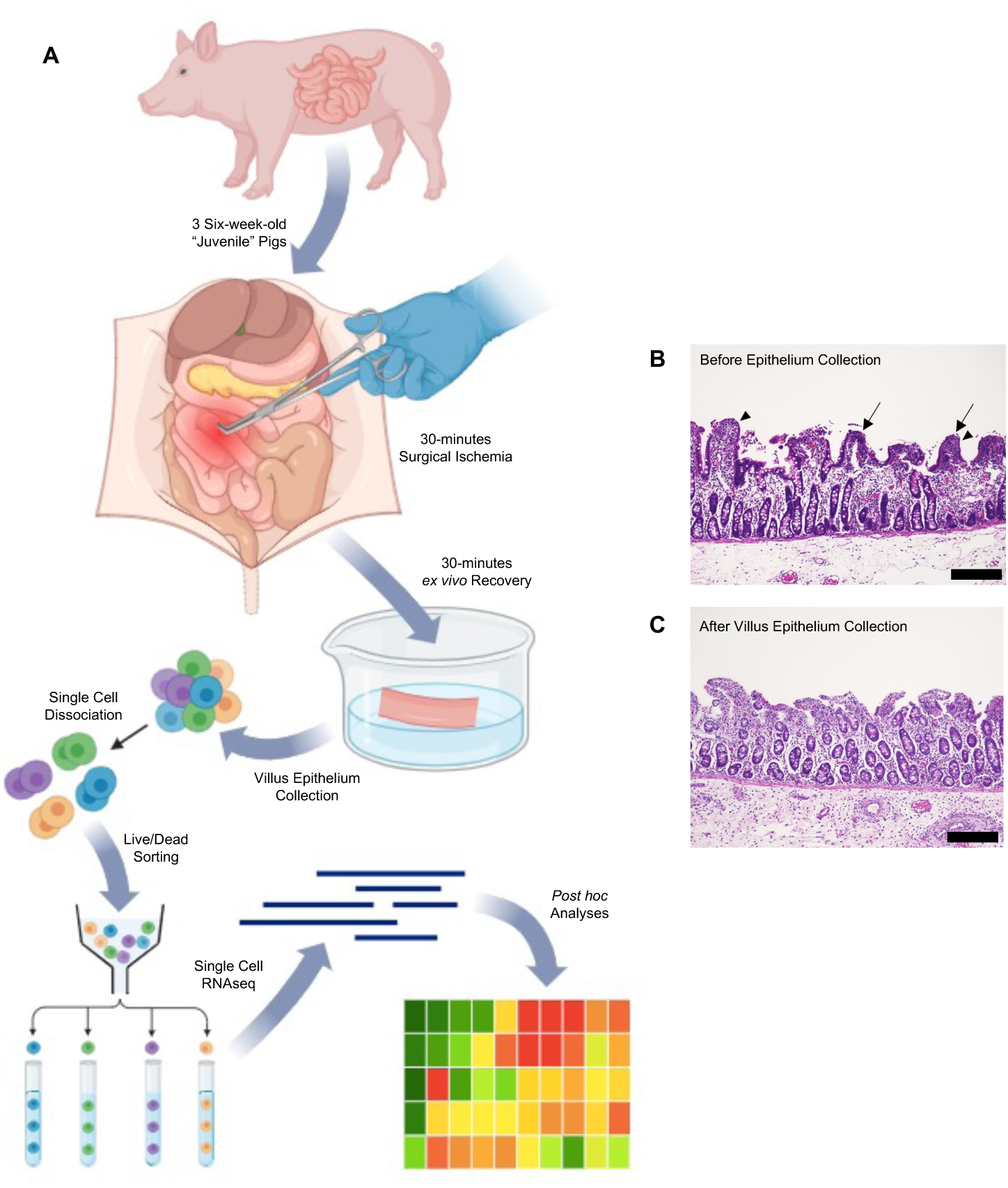
Ischemia-injured and *ex vivo* recovered jejunal mucosal tissues were prepared to produce a cell suspension enriched in restituting villus epithelial cells. **(A)** Illustrated workflow for single cell isolation approach. Segments of the distal jejunum of three commercial Yorkshire cross pigs were subjected to 30 minutes of ischemic injury followed by 30 minutes of *ex vivo* recovery. Upon sloughing and dissociation, cells isolated from the jejunum of all three pigs were pooled, sorted and sequenced. Created in Biorender. (B) Histomicrograph of ischemia-injured, *ex vivo* recovered jejunum prior to cell sloughing demonstrating restituting villus epithelial cells lining regions of mucosal denudation. (C) Histomicrograph of ischemia-injured, *ex vivo* recovered jejunum following cell sloughing reveals that this target population was captured and enriched for sequencing. Scale bar 100µm.

### IPEC-J2 cell scratch wound assay

IPEC-J2 cells an intestinal epithelial cell line originally isolated from the jejunum of a neonatal pig were cultured seeded onto Transwell inserts (Corning Cat# 3470) and maintained in DMEM/F12 supplemented with 10% heat-inactivated FBS, 5ng/mL insulin/ transferrin/ selenium supplement (Gibco Cat# 41400045) and 5 ng/mL recombinant human EGF (Gibco Cat# PHG0311).(22) Once visibly confluent, scratch-wounds were inflicted manually with a disposable 200µL micropipette tip. Wounds were imaged at 0-, 2-, 4- and 6-hours after scratch-wounding. Images were captured using a Keyence BZ-X800 All-in-One microscope (Osaka, Japan) using a 20x objective lens and images were processed using Keyence Software BZ Analyzer. Percent wound healing was calculated after measuring total wound area using ImageJ software (NIH, USA).

### Immunocytochemistry

To examine CSF-1 and CSF1R expression, IPEC-J2 cells were fixed at 8-hours of recovery from scratch wounding, permeabilized in 0.01% Triton-X PBS and incubated with rabbit anti-M-CSF polyclonal antibody (ThermoFisher PA542558, RRID:AB_260957) or mouse anti-CD115 monoclonal antibody (ThermoFisher MA528453, RRID:AB_2745415), in PBS. Cells were then incubated in goat anti-mouse conjugated to Alexa Fluor 488 (Thermo Fisher A28175; RRID:AB_2536161) or goat anti-rabbit conjugated to Alexa Fluor 647 (Thermo Fisher A21245; RRID:AB_2535813) diluted 1:500 for 2 hours at room temperature. Cell nuclei were stained DAPI (Invitrogen Cat# D1306) at a dilution of 1:1000 in PBS for 5 mins at room temperature. Images were captured using a Keyence BZ-X800 All-in-One microscope (Osaka, Japan) using a 20x objective lens and images were processed using Keyence Software BZ Analyzer.

## RESULTS

### Experimental protocols targeted restituting epithelial cells for sequencing

Previous experiments in our laboratory have demonstrated that the restituting cells active in post-ischemic intestinal healing are the superficial, villus epithelium lining the periphery of denuded mucosa. Therefore, we needed to ensure that our experimental methods had successfully isolated this target cell population for sequencing. A general schematic of jejunal epithelial cell procurement for cell sorting is provided (Fig 1A). Light microscopy of the ischemia-injured, *ex vivo* recovered jejunal segments prior to cell sloughing demonstrates the presence of restituting villus epithelial cells lining regions of mucosal denudation (Fig 1B, arrows). Light microscopy of jejunal segments following cell sloughing reveals that this target population is absent while the crypt epithelium remains (Fig 1C). Of the sloughed cells pooled from all three pigs, fluorescence-associated sorting identified and isolated 147,000 live epithelial cells based on negative staining for Annexin V and positive staining for EpCAM (Fig 2). Cells were sorted based upon doublet discrimination (Gates A and B), absence of Annexin V uptake (Gate C) and the presence of EpCAM uptake (Gate D). Of the first 100,000 cells sorted, there were 4,372 (Gate F) that passed Annexin V live/dead and EpCAM parameters during fluorescence-activated cell sorting. Therefore, our described protocols successful isolated the targeted restituting villus epithelial cells for sequencing.

**Figure 2.**
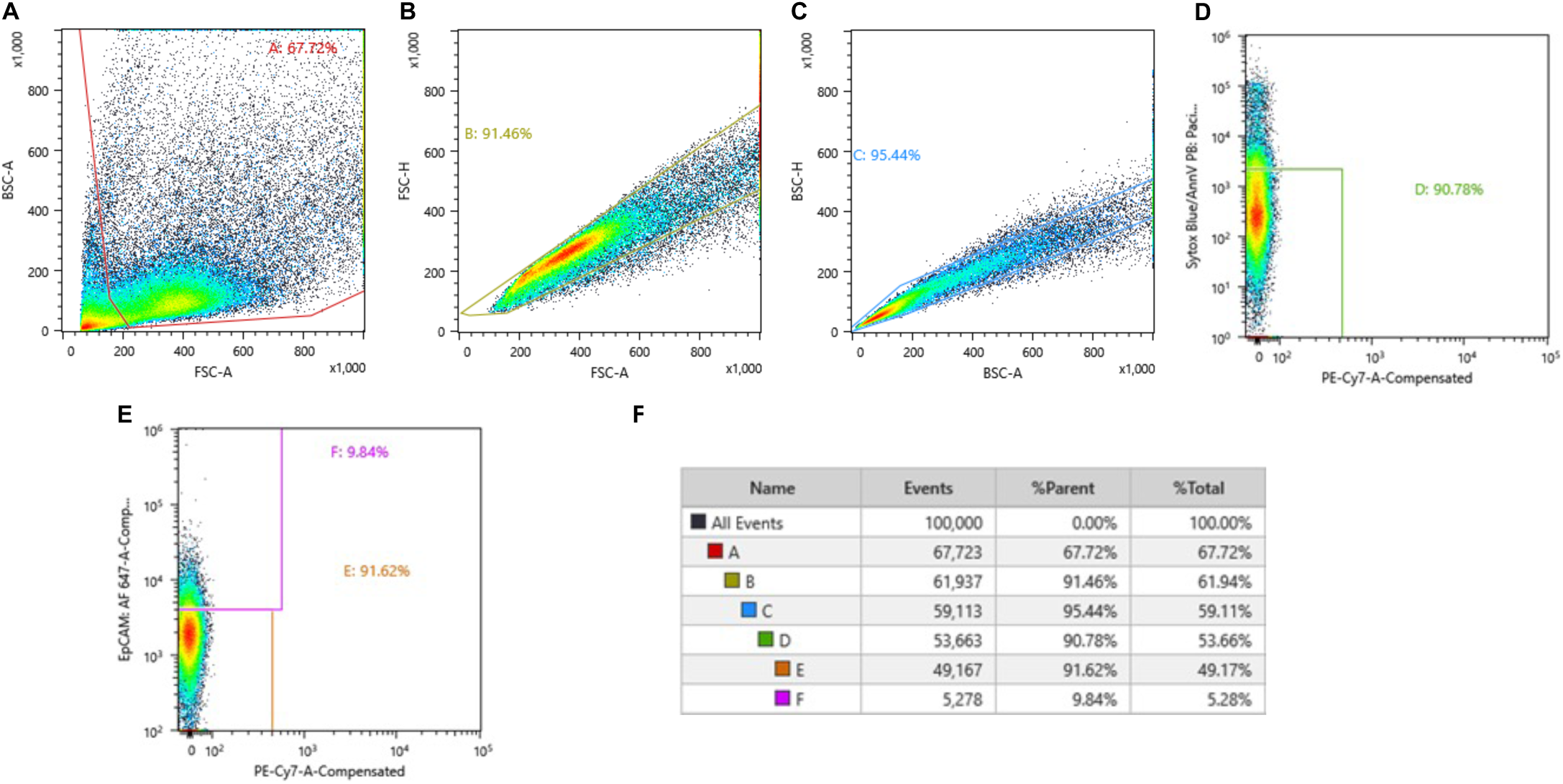
Single cells were sorted for to further enrich only single, live cells for sequencing. (A) All sorted cells graphed according to side and back scatter. (B and C) Cells were sorted based upon doublet discrimination using forward scatter (Gate A) and back scatter (Gate B). (D and E) Cells were further sorted based on absence of Annexin V uptake (Gate C) and the presence of EpCAM uptake (Gate D). (F) Of the first 100,000 cells sorted, there were 4,372 (Gate F) that passed Annexin V live/dead and EpCAM parameters during fluorescence-activated cell sorting.

### Clustering analysis revealed eight distinct intestinal epithelial cell populations

Single-cell transcriptomic profiling of these cells (n=1,359) revealed right distinct cell clusters. The percentages of the total cell population composed of each cluster were as follows: cluster 0 (36.72%), cluster 1 (24.80%), cluster 2 (21.92%), cluster 3 (5.52%), cluster 4 (3.31%), cluster 5 (2.87%), cluster 6 (2.58%), and cluster 7 (2.28%) (Fig 3A). Uniform Manifold Approximation and Projection (UMAP) highlights these eight cell clusters with some overlapping transcriptomic profiles shared between clusters 0 and 1 (Fig 3B-C). To infer the specific epithelial lineage represented by each cluster, each cluster was scored based on the expression of differentially expressed genes (DEGs) from lineages identified in a previous transcriptomic survey of adult adult human small intestine (Fig 3C).(13) Cluster 0 had the highest absorptive enterocyte score. Cluster 1 also had a high absorptive enterocyte score, although it scored highest for intestinal stem cells. Cluster 2 scored highest for intestinal stem cells. Cluster 3 scored highest for Tuft cells. Cluster 4 scored highest with enteroendocrine cells. Cluster 5 scored highest for follicle associated epithelium. Cluster 6 scored highest for transit amplifying cells and intestinal stem cells. Cluster 7 scored highest for Goblet cells and Best 4 cells. The numbers and percentages of the referenced lineage identity gene lists which were expressed in each cluster are provided in Tables 1 and 2, respectively.

**Figure 3.**
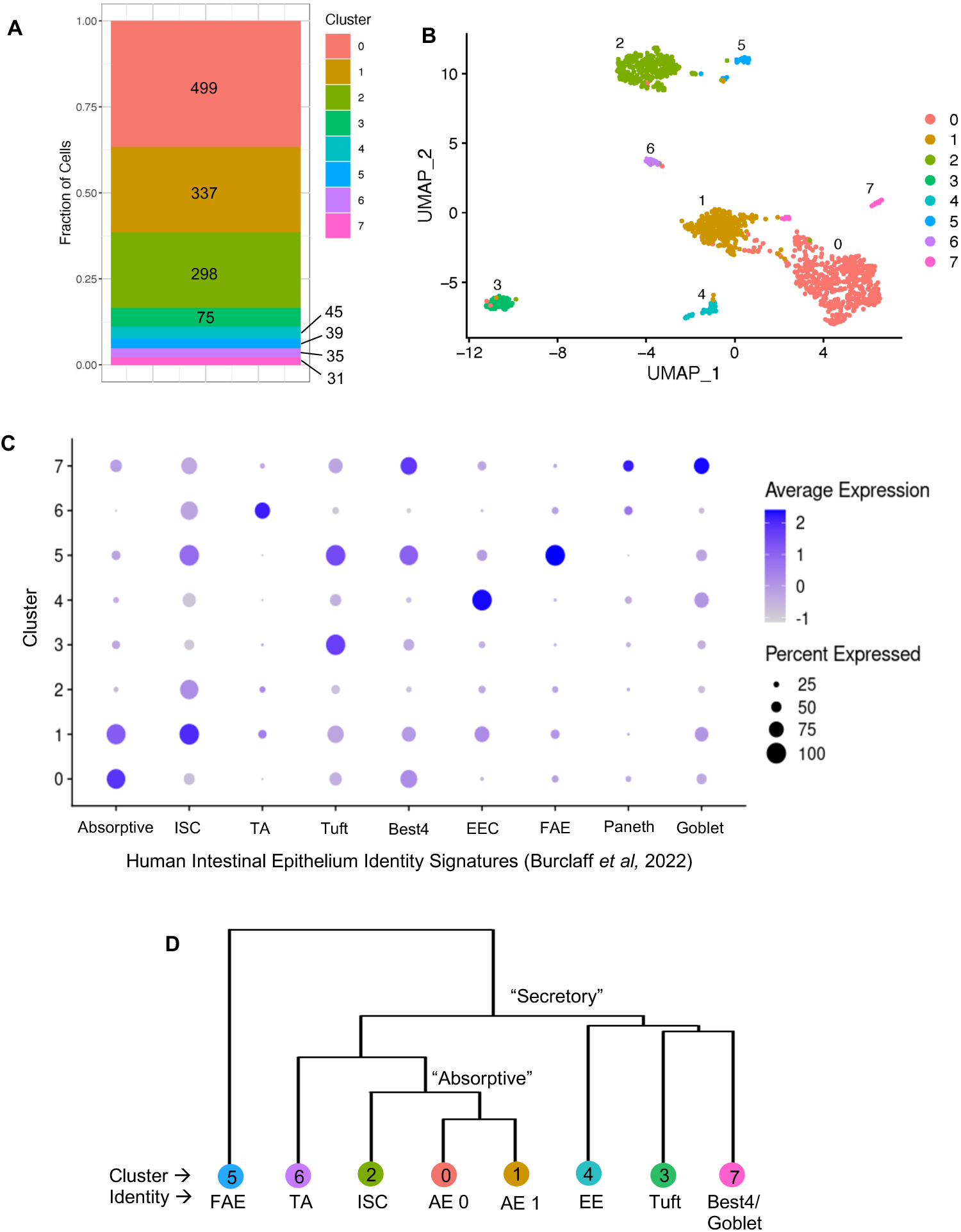
Clustering analysis isolated 8 cell clusters, which were putatively identified by transcriptomic profiling and hierarchical clustering. (A) The majority of the intestinal epithelial cells isolated from the three pigs and submitted for sequencing were assigned to clusters 0 and 1. Clusters 4, 5, 6 and 7 contained the fewest cells. (B) Unsupervised clustering demonstrated in this UMAP plot yielded 8 distinct clusters, and the largest clusters, 0 and 1, grouped closely and with some overlap. (C) The cell populations represented by the 8 clusters were presumptively identified by comparing each cluster’s transcriptomic profiles to those identified by Burclaff et al. (D) A simplicifolious clade dendrogram demonstrated a putative hierarchical relationship among the 8 clusters. Absorptive and the few proliferative lineages were loosely separated from secretory lineages in an early clade. Clusters 0 and 1 are more closely related to each other than any other cluster. AE= absorptive enterocyte, EE= enteroendocrine cell, FAE= follicle associated epithelial cell, ISC= intestinal stem cell and TA= transit amplifying cell.

**Table 1.** Number of genes from each identity’s gene list that are expressed in each cluster. AE= absorptive enterocyte, EE= enteroendocrine cell, FAE= follicle associated epithelial cell, ISC= intestinal stem cell and TA= transit amplifying cell.

| Identity | Genes in Identity List | Genes in Cluster |  |  |  |  |  |  |  |
| --- | --- | --- | --- | --- | --- | --- | --- | --- | --- |
|  |  | **0** | **1** | **2** | **3** | **4** | **5** | **6** | **7** |
| **AE** | 289 | 202 | 196 | 197 | 182 | 182 | 147 | 140 | 166 |
| **Best4** | 168 | 101 | 97 | 96 | 89 | 88 | 72 | 68 | 83 |
| **EEC** | 324 | 38 | 42 | 43 | 50 | 51 | 67 | 71 | 56 |
| **FAE** | 145 | 86 | 82 | 80 | 69 | 68 | 70 | 63 | 65 |
| **Goblet** | 276 | 175 | 171 | 173 | 159 | 164 | 139 | 127 | 165 |
| **ISC** | 67 | 41 | 40 | 40 | 36 | 38 | 29 | 33 | 38 |
| **Paneth** | 13 | 5 | 5 | 4 | 4 | 3 | 1 | 3 | 5 |
| **TA** | 358 | 279 | 279 | 280 | 280 | 281 | 283 | 281 | 279 |
| **Tuft** | 957 | 377 | 377 | 376 | 376 | 375 | 373 | 375 | 377 |

**Table 2.** Percentage of genes from each identity’s gene list that that are expressed in each cluster. AE= absorptive enterocyte, EE= enteroendocrine cell, FAE= follicle associated epithelial cell, ISC= intestinal stem cell and TA= transit amplifying cell.

| Identity | Percent Expressed in Cluster |  |  |  |  |  |  |  |
| --- | --- | --- | --- | --- | --- | --- | --- | --- |
|  | 0 | 1 | 2 | 3 | 4 | 5 | 6 | 7 |
| **AE** | 69.9 | 67.8 | 68.2 | 63.0 | 63.0 | 50.9 | 48.4 | 57.4 |
| **Best4** | 60.1 | 57.7 | 57.1 | 53.0 | 52.4 | 42.9 | 40.5 | 49.4 |
| **EEC** | 11.7 | 13.0 | 13.3 | 15.4 | 15.7 | 20.7 | 21.9 | 17.3 |
| **FAE** | 59.3 | 56.6 | 55.2 | 47.6 | 46.9 | 48.3 | 43.4 | 44.8 |
| **Goblet** | 63.4 | 62.0 | 62.7 | 57.6 | 59.4 | 50.4 | 46.0 | 59.8 |
| **ISC** | 61.2 | 59.7 | 59.7 | 53.7 | 56.7 | 43.3 | 49.3 | 56.7 |
| **Paneth** | 38.5 | 38.5 | 30.8 | 30.8 | 23.1 | 7.7 | 23.1 | 38.5 |
| **TA** | 77.9 | 77.9 | 78.2 | 78.2 | 78.5 | 79.1 | 78.5 | 77.9 |
| **Tuft** | 39.4 | 39.4 | 39.3 | 39.3 | 39.2 | 39.0 | 39.2 | 39.4 |

To further support this putative cluster identification, a dendrogram was constructed to explore the hierarchal relationships among each cluster (Fig 3D). A simplicifolious clade separated Cluster 5, the presumed follicle associated epithelial cells, from all other clusters. The remaining 7 clusters divided between two clades; clusters 6, 2, 0 and 1 organized into an “absorptive” clade while clusters 4, 3 and 7 were organized into a “secretory” clade. The “absorptive” clade was further organized as clusters 0 and 1, the presumed absorptive enterocytes, being the most similar transcriptomically followed by cluster 2, the presumed intestinal stem cells, and then cluster 6, the presumed intestinal stem cells / transit amplifying cells. Within the “secretory” clade, clusters 3 and 7, the presumed Tuft cells and the presumed Best4/Goblet cells respectively, were the most similar transcriptomically followed by cluster 4, the presumed enteroendocrine cells. The constellation of the UMAP, dendrogram and dot plot provided strong support for our putative cluster identification.

### Cluster 0 demonstrates relatively increased expression of physiologic cellular migration pathways and highlights potential key mediators of restitution

Given the results of unsupervised clustering, Clusters 0 and 1, which make up most of the sequenced cell population, are presumed to be two sub-phenotypes of absorptive enterocytes, and we hypothesized that one of these two is a subset of cells that are diverging into the restitution phenotype. To identify this restitution cluster, we performed comparative analysis to discover genes that specifically define the differences between these two discrete clusters of absorptive enterocytes. IPA^®^ was used to investigate which of these two clusters demonstrated increased relative expression of physiologic pathways related to cell movement. Several key diseases and functions were enriched in Cluster 0 relative to Cluster 1, including Gastrointestinal Disease, Cell Death and Survival, Cell-to-Cell Signaling and Interaction, Cellular Movement, Cell Morphology and Cellular Assembly and Organization, each of which have direct relevance to epithelial restitution responses (Fig 4A). The most relevant Disease and Process, Cell Movement, was further examined by plotting a heat map for each pathway within that group. This more granular analysis demonstrates that all the differentially expressed Cellular Movement pathways are enriched in cluster 0 relative to cluster 1, including “Migration of Cells” (Fig 4B). These data lead us to surmise that cluster 0 represents a sub-population of absorptive enterocytes which are undergoing transcriptional responses to shift into the restitution phenotype and repair the damaged epithelial barrier.

**Figure 4.**
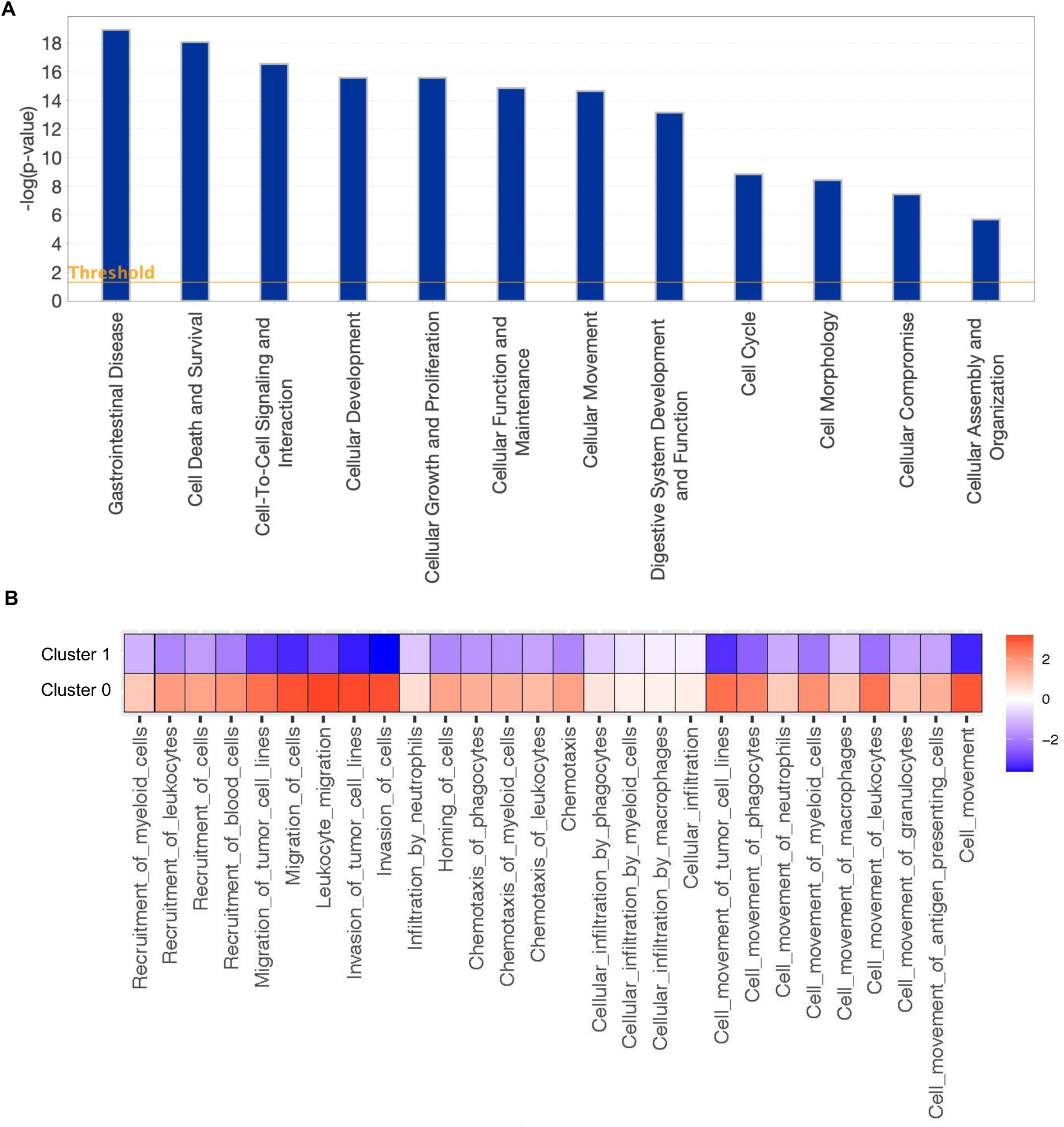
DEGs in cluster 0 versus cluster 1 demonstrate that Cluster 0 is the restituting subset of absorptive enterocytes. (A) Diseases and Functions groups which are differentially represented in cluster 0 relative to cluster 1 include responses highly relevant to enterocyte restitution. (B) Numerous specific pathways within the “Cell Movement” group are upregulated in cluster 0.

### Upstream Regulators analysis predicts cluster 0 transcriptional responses are regulated by Colony-Stimulating Factor-1 (CSF-1)

Knowing that cluster 0 is the subset of absorptive enterocytes which are shifting into restitution programs during epithelial barrier repair, we sought to predict which signaling molecules may be driving these transcriptional changes. The top five predicted upstream regulators (predictions with the smallest p-values) included tumor necrosis factor (TNF), interferon γ (IFNG), interleukin 1β (IL1B), colony stimulating factor-1 (CSF1) and transforming growth factor β1 (TGFB1) (Fig 5A). As our interest in these upstream regulators was borne from the age-dependent restitution response observed in neonatal piglets which can be rescued by direct application of mucosal homogenates from ischemia-injured jejunum of juvenile pigs, but not neonatal pigs, we sought to define which of these predicted regulators’ expression is induced by ischemia in an age-dependent manner.(11) To do this, bulk transcriptional sequencing data from those experiments, which are publicly available in the Gene Expression Omnibus (https://www.ncbi.nlm.nih.gov/geo/, accession #GSE212533), was used to generate a heat map of total mRNA counts for each predicted regulator in the homogenates of neonatal and juvenile control and ischemic jejunal mucosa.(12) This revealed that the only regulator demonstrating ischemia-induced mRNA expression in juvenile jejunum, but not neonatal jejunum, is *CSF1* (Fig 5B). The mechanistic network in cluster 0 which predicted its upstream regulation by CSF-1 includes 153 genes (Fig 5A). To validate that this set of CSF-1-responsive genes in cluster 0 is regulating processes directly related to specific restitution responses in this cluster, these 153 genes were tested for Gene Ontology (GO) and many pathways distinctive of epithelial restitution responses following ischemic injury were identified (Fig 5C). For example, “negative regulation of cell-cell adhesion”, “response to hypoxia” and “positive regulation of cell migration” stood out. Additionally, a subset of genes expressed in cluster 0 known to respond to CSF-1 and that may play a role in cellular movement functions. (Fig. 5D).

**Figure 5.**
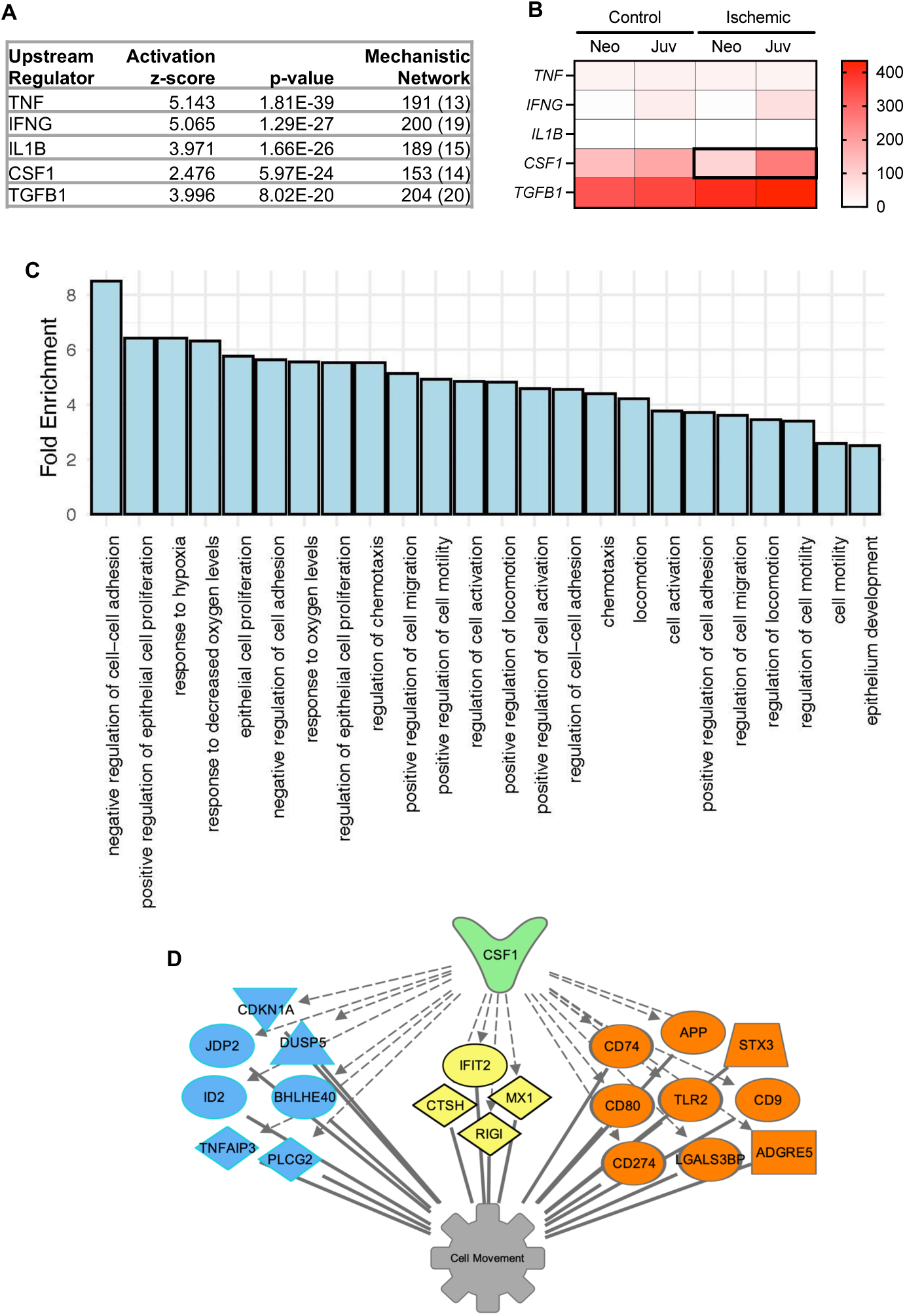
upstream regulators analysis highlights CSF-1 as a potential age-dependent regulator of interest. (A) Predicted upstream regulators of cluster 0 phenotype are routinely implicated in intestinal barrier maintenance, except for one novel regulator, colony-stimulating factor-1 (CSF-1). (B) Of the top-5 predicted upstream regulators, total mRNA detected in homogenized mucosa reveals age-dependent ischemia-induced upregulation of only CSF1 (N=3). (C) GO Enrichment Analysis for 153 CSF-1-responsive DEG expressed in cluster 0. (D) My Pathways tool identified a subset of DEG in Cluster 0 which encode nuclear (blue), cytoplasmic (yellow) or membrane bound (orange) signaling proteins known to respond to CSF-1 Predicted relationships between CSF-1 and these genes (hashed grey arrows) and these genes and the down-stream function “Cell movement” (solid grey lines) are represented graphically.

### Interrogation of CSF-1 signaling in our pig intestinal injury model offers preliminary proof-of-concept for its role in age-dependent regulation of restitution

Given the prediction that CSF-1 is an upstream regulator of restitution responses in juvenile enterocytes, and *CSF1* is induced by ischemia in an age-dependent manner in mucosal homogenates, we sought to examine age-dependent mucosal expression patterns of this cytokine and its receptor, colony-stimulating factor-1 receptor (CSF1R) during injury and repair. Histological slides from experiments previously reported by our group were stained for CSF-1 and CSF1R.(11) In acutely ischemia-injured mucosa, expression of CSF1R was observed in wound-adjacent epithelial cells regardless of age, but CSF-1 co-localized with CSF1R only in juvenile wound-adjacent epithelial cells (Fig 6A). Similarly, injured mucosa that has undergone *ex vivo* recovery also demonstrated expression of CSF1R in the wound-adjacent epithelium regardless of age, and CSF-1 co-localization with epithelial CSF1R was detected in juveniles but not neonates (Fig. 6B). However, neonatal mucosa which was treated with direct application of juvenile homogenized mucosa during *ex vivo* recovery demonstrated CSF-1 co-localization with epithelial CSF1R in the “rescued” restituting epithelium (Fig. 6B). These findings validate the potential role of CSF-1 signaling in mediating restitution in juveniles and in “rescued” neonates. To further validate a mechanistic link between CSF-1 signaling and epithelial restitution, the effects of CSF1R inhibition was tested *in vitro*. Firstly, IPEC-J2 cells were confirmed to express both CSF-1 and CSF1R during scratch wound restitution *in vitro* (Fig. 6C). Finally, CSF1R antagonist BLZ945 inhibits restitution of IPEC-J2 scratch wounds *in vitro* in a dose-dependent manner (P=0.0081 by one-way ANOVA) suggesting that CSF-1 signaling is required for efficient restitution of porcine intestinal epithelial cells (Fig. 6D).

**Figure 6.**
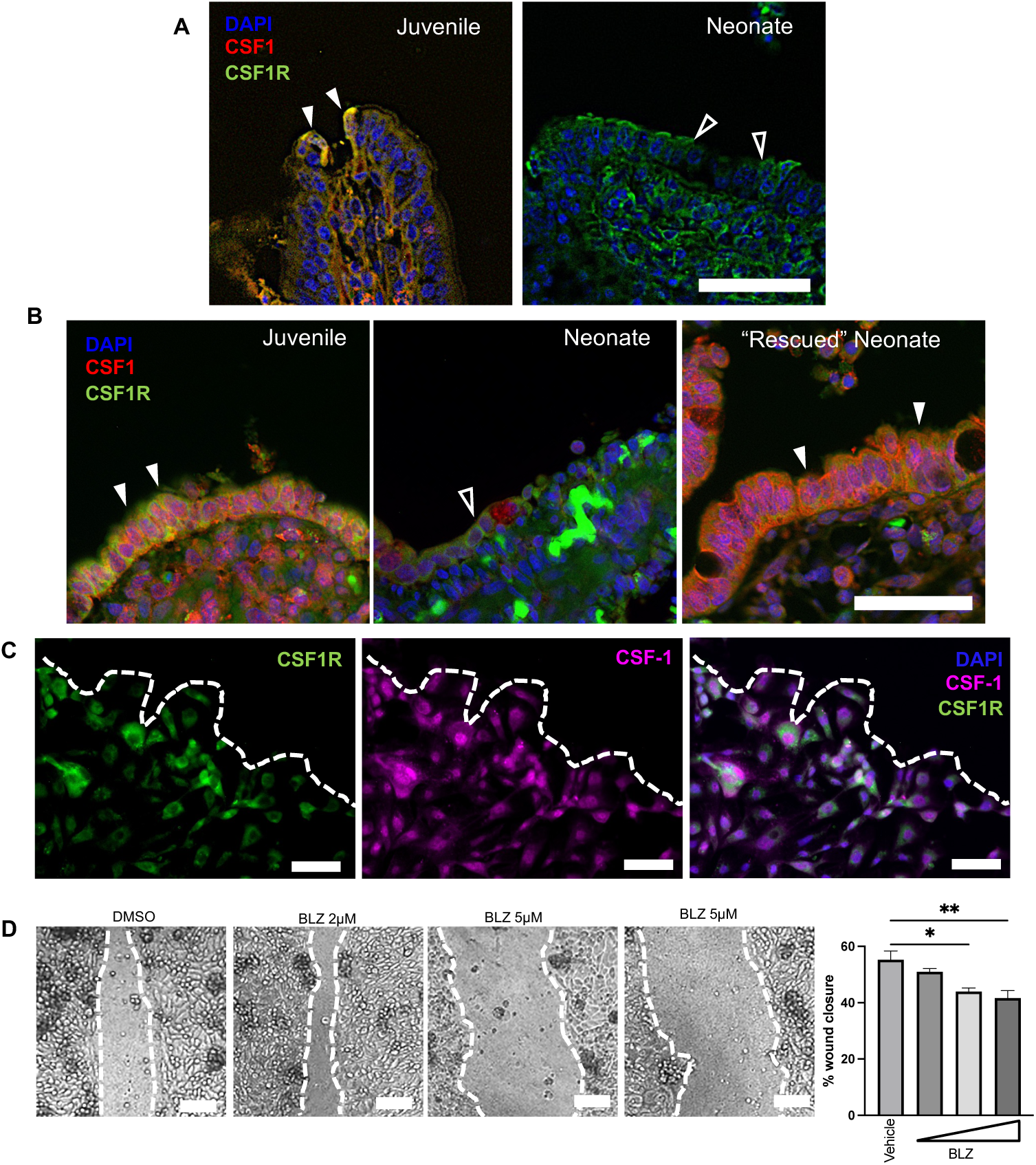
Interrogation of CSF-1 signaling in our pig intestinal injury model offers positive proof-of-concept for its role in age-dependent regulation of restitution. (A) Representative IHC images of ischemia-injured mucosa validates expression of CSF1R (green) in wound-adjacent epithelial cells regardless of age (closed and open arrowheads), but CSF-1 (red) co-localization with CSF1R is detected only in juvenile wound-adjacent epithelial cells (closed arrowheads). (B) Representative IHC images of injured mucosa after *ex vivo* recovery reveals expression of CSF1R (green) in epithelium regardless of age, but CSF-1 (red) co-localization with epithelial CSF1R is detected only in juveniles and in neonates rescued with direct application of juvenile homogenized mucosa during recovery (closed arrowheads). (C) IPEC-J2 express CSF-1 and CSF1R during scratch wound restitution *in vitro*. Nuclei labeled with DAPI (blue). Scale bar 50µm. (D) CSF1R antagonist BLZ inhibits restitution of IPEC-J2 scratch wounds *in vitro* in a dose-dependent manner. N=3. P=0.0081 by one-way ANOVA. *P≤0.05, **P≤0.01 by Dunnett’s multiple comparisons test. Scale bar 100µm.

## DISCUSSION

Initial analysis of the transcriptomic profiles of all sloughed cells revealed the presence of 8 distinct cell clusters. Clusters 0 and 1 were easily identifiable as absorptive enterocytes based on the average gene expression and percent of cells expressing genes associated with this lineage. Taking the identification of these two clusters into account, the fact that absorptive enterocytes account for 61.5% of the sloughed, sequenced epithelial cells that passed quality control (Fig 3A) validates that the epithelial sloughing protocol successfully enriches the superficial villus epithelium. The next largest cluster, cluster 2, demonstrated greatest overlap with ISC, so these may represent small numbers of ISC obtained from the mucosal crypts. DEGs used to identify cell clusters were assigned to human cell lineages, therefore it is possible that slight transcriptomic differences between human and pig intestinal stem cells account for a slight misidentification of this cluster. For example, it is possible that cluster 2 in fact represents pig transit amplifying cells, whose position along the crypt-villus axis makes it more plausible over ISC that they would be included in this villus-enriched sample population. Further, the identification method attempted here was limited by the collection of villus-enriched samples, thus there was not a complete profile of every epithelial subtype present in the pooled sample, reducing the cellular diversity for initial unbiased clustering which would be expected to reduce some of the differential clustering resolution as compared to the more complete surveys acheived by Burclaff *et al* (13). However, the calculated hierarchical relationships among the 8 clusters (Fig 3D), supports the lineage identifications suggested by overlap with the human dataset. Cluster 5, the suspected follicle associated epithelium, is separated from the other 7 clusters into its own clade. Follicle associated epithelial cells, which are also referred to as M cells, are suspected to originate from intestinal stem cells that are anatomically distinct from those that differentiate into absorptive and secretory enterocytes(23). Therefore, it is unsurprising that this cluster is organized into a simplicifolious clade separate from all other clusters. Following the branching off of cluster 5, the next branching separates clusters 6, 2, 0 and 1 from clusters 4, 3 and 7. This again corroborates identification of each cluster because this branching distinguishes the absorptive lineages, namely the absorptive enterocytes of clusters 0 and 1, from the secretory enterocytes, namely the enteroendocrine cells (cluster 4), the tuft cells (cluster 3), and the Best4/Goblet cells (cluster 7). In addition, as represented by the shortest branches within the dendrogram, the two subpopulations of absorptive enterocytes (clusters 0 and 1) are the most closely related with respect to their transcriptomic profiles. Finally, this dendrogram’s hierarchical organization of the isolated clusters closely mimics dendrograms and trajectory analyses of intestinal epithelial lineages described within separate studies(13, 24). Certainly, the identity of clusters 2-7 are not definitive within the scope of the present report.

Given that the clusters 0 and 1 are both absorptive enterocyte lineages and are very closely related as demonstrated by unsupervised and hierarchal clustering, we surmised that one of these populations is a subtype of absorptive enterocytes which is altering its transcriptional programs in response to ischemic injury and diverging from the other cluster of absorptive enterocytes into a restitution subcluster. Indeed, DEG between clusters 0 and 1 reveal upregulation of Diseases and Processes highly relevant to restitution processes in cluster 0 relative to cluster 1, so it may be concluded that cluster 0 represents the population of restituting enterocytes (Fig 4A). It can also be noted that differential expression of the Diseases and Processes “Digestive System Development and Function” and “Gastrointestinal Disease” between these clusters further validates the accuracy of sequencing study; these profiles represent expected transcriptomic profile signatures of gastrointestinal epithelium and, in particular, our surgical injury model inflicting a state of gastrointestinal disease. Upregulation of “Cell Death and Survival” and “Cellular Compromise” functions substantiates that these cells are undergoing signaling responses key to their survival after severe cellular stress and injury, so that they may participate in mucosal repair. “Cell morphology” and “Cellular Assembly and Organization” and “Cellular Movement” functions indicates a clear alteration in cell functions in Cluster 0 as they relate to cellular depolarization, detachment, and migration – processes critical for the restitution cellular phenotype.(25) Indeed, when further interrogating the numerous transcriptional pathways which define the larger “Cellular Movement” function, we can drill down to observe cluster 0 enrichment of the specific pathways “migration of cells” “invasion of cells” and “homing of cells” which indicate that these cells are undergoing directed cell crawling programs related to restitution (Fig 4B).

These findings indicate that single cell transcriptomics has successfully defined a distinct population of intestinal enterocytes undergoing a swift phenotypic transition into a wound-healing cellular program in this unique pig model. Others have applied a similar approach skin wound healing, and these studies have revealed important findings in the context of more complex, stem-cell based subacute wound healing in injured dermal tissues.(26) These studies provide valuable insights as to the roles of diverse cell types in complex stratified squamous epithelial wound healing. The finely focused set of data generated by the present study allows for precise interrogation of the enterocyte reprogramming which occurs in a matter of minutes following a very carefully controlled loss of columnar epithelium by ischemic injury. These data present a complete transcriptomic picture of the very specific, speedy response by wound-adjacent enterocytes which are depolarizing and initiating directed migration across the mucosal erosion they are sensing before the proliferative subacute phase of repair is initiated by ISC.(27) This set of DEG defines the complete, precise and distinct transcriptional changes that are occurring in the enterocytes which are depolarizing and crawling as compared to adjacent absorptive enterocytes which are not undergoing this change. Presently, we have identified an age-dependent absence of this restitution program in our neonatal pig model of intestinal injury.(11) While neonatal restitution is inducible with direct application of the intestinal milieu from restitution-competent juvenile aged animals, we do not know which signals are present in this milieu which are responsible for inducing restitution in neonates. Given that we have captured this unique signature of DEG, we applied the powerful *post hoc* analyses in IPA^®^ which allow prediction of upstream regulators of discrete transcriptomic signatures in order to predict which signaling molecules might be key contributors to this critical phenotypic change.(28) Upstream regulators analysis predicted that the transcriptional changes in Cluster 0 may be driven by several potential molecules including TNF, IFN-γ, IL-1β, CSF-1 and TGF-β1 (Fig 5A). TNF, IFN-γ, IL-1β, and TGF-β1 are well-known to regulate barrier function and inflammatory processes in the intestine, and thus serve to validate this approach to predict regulators of mucosal injury responses.(29–37) However, in the context of our age-dependent repair model, we examined these candidates’ transcriptional expression in the banked homogenized mucosal tissues from our previously-reported study(11) and found ischemia-induction in juveniles, but not neonates, of only *CSF-1* (Fig 5B). Thus, CSF-1 could be an important target for ongoing study, as it is the only predicted regulator transcriptionally induced in juvenile, but not neonatal, jejunum in response to injury which corresponds with the age-dependency of mucosal repair.

CSF-1 is a cytokine produced by many diverse cell populations primarily implicated in macrophage maintenance, but which is known to also regulate epithelial cells in several contexts, including recent report of it inducing migratory pathways in an epithelial cell line.(38–43) Validating of a potential role for CSF-1 signaling in regulating induction restitution in our model, we found juvenile mucosa demonstrates strong co-localization of CSF1 and CSF-1 receptor (CSF1R) very rapidly following ischemic injury in wound-adjacent epithelial cells and continued co-localization in the restituted epithelium (Fig 6 A and B). Further, we have confirmed CSF1R expression in injured neonatal epithelium and co-localization of CSF1 and CSF1R in restituted neonatal epithelium following its rescue with the juvenile homogenized mucosa (Fig 6B). Moreover, *in vitro* experiments presented here demonstrate that CSF1R antagonist BLZ945 inhibits restitution of scratch wounds in porcine jejunal epithelial monolayers in a dose dependent manner (Fig 6 D). These data serve to support the premise that CSF-1 signaling is an important candidate for ongoing study in the context of age-dependent restitution responses in the small intestine and validate this single cell transcriptomic approach for predicting upstream regulators of very precise, acute cellular reprogramming responses in the intestinal epithelium.

## CONCLUSIONS

The current study validates an approach to target and isolate acutely restituting intestinal epithelial cells for high resolution transcriptomic profiling. Analyzing this target cell population through single cell resolution provides powerful insight into the transcriptomics that define this transient restituting enterocyte phenotype. These predictions will inform future work to delve deeper into the mechanics of the restituting phenotype by investigating the role of predicted upstream regulators, such as CSF-1. This has the potential to inform novel treatment modalities to promote restitution and barrier recovery in patients with severe intestinal injury. Furthermore, we hope that these sequencing data demonstrate an approach toward understanding the pathophysiology of cellular injury and repair beyond acute epithelial restitution to guide advancements in precision medicine.

## DATA AVAILABILITY

Source data for this study are openly available at GEO Accession number GSE271071 (https://www.ncbi.nlm.nih.gov/geo/query/acc.cgi?acc=GSE271071).

## ACKNOWLEDGEMENTS

We would like to thank Tiffany Pridgen and Courtney Deck for assisting in lab management and lab bench work, North Carolina State University’s Lab Animal Resources and Central Procedures Lab for facilitating our experimental animal model and surgical procedures.

## AUTHOR CONTRIBUTIONS

Conceived and designed research (ECR, ALZ, STM). Performed experiments (ECR, ALZ). Analyzed data (ECR, ALZ, JMS, IGM). Interpreted results of experiments (ECR, ALZ, STM, JMS, IGM, JO, ATB). Prepared figures (ECR, ALZ, JMS, IGM). Drafted manuscript (ECR, ALZ). Edited and revised manuscript (ECR, ALZ, STM, JMS, IGM, JO, ATB). Approved final version of manuscript (ECR, ALZ).

---

**Supplemental Figure 1. Secondary antibody control immunofluorescent histology images**. Scale Bar 300µm.

## BIBLIOGRAPHY

1. Paclik D, Lohse K, Wiedenmann B, Dignass AU, and Sturm A. Galectin-2 and −4, but not galectin-1, promote intestinal epithelial wound healing in vitro through a TGF-beta-independent mechanism. Inflammatory bowel diseases 14: 1366–1372, 2008.

2. Nusrat A, Delp C, and Madara JL. Intestinal epithelial restitution. Characterization of a cell culture model and mapping of cytoskeletal elements in migrating cells. Journal of Clinical Investigation 89: 1501–1511, 1992.

3. Podolsky DK. Healing the epithelium: solving the problem from two sides. Journal of gastroenterology 32: 122–126, 1997.

4. Jacobi SK, Moeser AJ, Corl BA, Harrell RJ, Blikslager AT, and Odle J. Dietary long-chain PUFA enhance acute repair of ischemia-injured intestine of suckling pigs. The Journal of nutrition 142: 1266–1271, 2012.

5. Moeser AJ, Prashant K. Nighot, Kathleen A. Ryan, Jenna G. Wooten, and Anthony T. Blikslager. Prostaglandin-mediated inhibition of Na+/H+ exchanger isoform 2 stimulates recovery of barrier function in ischemia-injured intestine. American journal of physiology Gastrointestinal and liver physiology 291: G885–G894, 2006.

6. Ziegler A, Gonzalez L, and Blikslager A. Large Animal Models: The Key to Translational Discovery in Digestive Disease Research. Cell Mol Gastroenterol Hepatol 2: 716–724, 2016.

7. Gonzalez LM, Moeser AJ, and Blikslager AT. Animal models of ischemia-reperfusion-induced intestinal injury: progress and promise for translational research. American journal of physiology Gastrointestinal and liver physiology 308: G63–75, 2015.

8. Roura E KS, Lallès JP, Le Huerou-Luron, de Jager N, Schuurman T, Val-Laillet D. Critical review evaluating the pig as a model for human nutritional physiology. Nutritional Reseach Reviews 29: 60–90, 2016.

9. Swindle MM, Makin A, Herron AJ, Clubb FJ, Jr., and Frazier KS. Swine as models in biomedical research and toxicology testing. Veterinary pathology 49: 344–356, 2012.

10. Burrin D, Sangild PT, Stoll B, Thymann T, Buddington R, Marini J, Olutoye O, and Shulman RJ. Translational Advances in Pediatric Nutrition and Gastroenterology: New Insights from Pig Models. Annu Rev Anim Biosci 8: 321–354, 2020.

11. Ziegler AL, Pridgen TA, Mills JK, Gonzalez LM, Van Landeghem L, Odle J, and Blikslager AT. Epithelial restitution defect in neonatal jejunum is rescued by juvenile mucosal homogenate in a pig model of intestinal ischemic injury and repair. PLoS One 13: e0200674, 2018.

12. Ziegler AL, Caldwell ML, Craig SE, Hellstrom EA, Sheridan AE, Touvron MS, Pridgen TA, Magness ST, Odle J, Van Landeghem L, and Blikslager AT. Enteric glial cell network function is required for epithelial barrier restitution following intestinal ischemic injury in the early postnatal period. American journal of physiology Gastrointestinal and liver physiology 326: G228–G246, 2024.

13. Burclaff J, Bliton RJ, Breau KA, Ok MT, Gomez-Martinez I, Ranek JS, Bhatt AP, Purvis JE, Woosley JT, and Magness ST. A Proximal-to-Distal Survey of Healthy Adult Human Small Intestine and Colon Epithelium by Single-Cell Transcriptomics. Cellular and Molecular Gastroenterology and Hepatology 13: 1554–1589, 2022.

14. Wiarda JE, Becker SR, Sivasankaran SK, and Loving CL. Regional epithelial cell diversity in the small intestine of pigs. J Anim Sci 101: 2023.

15. Gao S, Yan L, Wang R, Li J, Yong J, Zhou X, Wei Y, Wu X, Wang X, Fan X, Yan J, Zhi X, Gao Y, Guo H, Jin X, Wang W, Mao Y, Wang F, Wen L, Fu W, Ge H, Qiao J, and Tang F. Tracing the temporal-spatial transcriptome landscapes of the human fetal digestive tract using single-cell RNA-sequencing. Nature cell biology 20: 721–734, 2018.

16. Wang Y, DiSalvo M, Gunasekara DB, Dutton J, Proctor A, Lebhar MS, Williamson IA, Speer J, Howard RL, Smiddy NM, Bultman SJ, Sims CE, Magness ST, and Allbritton NL. Self-renewing Monolayer of Primary Colonic or Rectal Epithelial Cells. Cell Mol Gastroenterol Hepatol 4: 165–182 e167, 2017.

17. Srivastava RK, Singh RB, Pujarula VL, Bollam S, Pusuluri M, Chellapilla TS, Yadav RS, and Gupta R. Genome-Wide Association Studies and Genomic Selection in Pearl Millet: Advances and Prospects. Front Genet 10: 1389, 2019.

18. Soneson C, Matthes KL, Nowicka M, Law CW, and Robinson MD. Isoform prefiltering improves performance of count-based methods for analysis of differential transcript usage. Genome biology 17: 12, 2016.

19. Hippen AA, Falco MM, Weber LM, Erkan EP, Zhang K, Doherty JA, Vaharautio A, Greene CS, and Hicks SC. miQC: An adaptive probabilistic framework for quality control of single-cell RNA-sequencing data. PLoS Comput Biol 17: e1009290, 2021.

20. Kolberg L, Raudvere U, Kuzmin I, Vilo J, and Peterson H. gprofiler2 -- an R package for gene list functional enrichment analysis and namespace conversion toolset g:Profiler. F1000Res 9: 2020.

21. Satija R, Farrell JA, Gennert D, Schier AF, and Regev A. Spatial reconstruction of single-cell gene expression data. Nat Biotechnol 33: 495–502, 2015.

22. Schierack P, Nordhoff M, Pollmann M, Weyrauch KD, Amasheh S, Lodemann U, Jores J, Tachu B, Kleta S, Blikslager A, Tedin K, and Wieler LH. Characterization of a porcine intestinal epithelial cell line for in vitro studies of microbial pathogenesis in swine. Histochemistry and cell biology 125: 293–305, 2006.

23. Corr SC, Gahan CC, and Hill C. M-cells: origin, morphology and role in mucosal immunity and microbial pathogenesis. FEMS Immunol Med Microbiol 52: 2–12, 2008.

24. Gehart H, van Es JH, Hamer K, Beumer J, Kretzschmar K, Dekkers JF, Rios A, and Clevers H. Identification of Enteroendocrine Regulators by Real-Time Single-Cell Differentiation Mapping. Cell 176: 1158–1173 e1116, 2019.

25. Rankin CR, Hilgarth RS, Leoni G, Kwon M, Den Beste KA, Parkos CA, and Nusrat A. Annexin A2 regulates beta1 integrin internalization and intestinal epithelial cell migration. The Journal of biological chemistry 288: 15229–15239, 2013.

26. Joost S, Jacob T, Sun X, Annusver K, La Manno G, Sur I, and Kasper M. Single-Cell Transcriptomics of Traced Epidermal and Hair Follicle Stem Cells Reveals Rapid Adaptations during Wound Healing. Cell Rep 25: 585–597 e587, 2018.

27. Stewart AS, Schaaf CR, Luff JA, Freund JM, Becker TC, Tufts SR, Robertson JB, and Gonzalez LM. HOPX(+) injury-resistant intestinal stem cells drive epithelial recovery after severe intestinal ischemia. American journal of physiology Gastrointestinal and liver physiology 321: G588–G602, 2021.

28. Kramer A, Green J, Pollard J, Jr., and Tugendreich S. Causal analysis approaches in Ingenuity Pathway Analysis. Bioinformatics 30: 523–530, 2014.

29. Billmeier U, Dieterich W, Neurath MF, and Atreya R. Molecular mechanism of action of anti-tumor necrosis factor antibodies in inflammatory bowel diseases. World journal of gastroenterology 22: 9300–9313, 2016.

30. Dignass AU, and Podolsky DK. Cytokine modulation of intestinal epithelial cell restitution: central role of transforming growth factor beta. Gastroenterology 105: 1323–1332, 1993.

31. Bruewer M, Utech M, Ivanov AI, Hopkins AM, Parkos CA, and Nusrat A. Interferon-gamma induces internalization of epithelial tight junction proteins via a macropinocytosis-like process. FASEB J 19: 923–933, 2005.

32. Liu H, Wang P, Cao M, Li M, and Wang F. Protective role of oligomycin against intestinal epithelial barrier dysfunction caused by IFN-gamma and TNF-alpha. Cellular physiology and biochemistry : international journal of experimental cellular physiology, biochemistry, and pharmacology 29: 799–808, 2012.

33. Yang S, Yu M, Sun L, Xiao W, Yang X, Sun L, Zhang C, Ma Y, Yang H, Liu Y, Lu D, Teitelbaum DH, and Yang H. Interferon-gamma-induced intestinal epithelial barrier dysfunction by NF-kappaB/HIF-1alpha pathway. Journal of interferon & cytokine research : the official journal of the International Society for Interferon and Cytokine Research 34: 195–203, 2014.

34. Kretzschmar K, and Clevers H. IFN-gamma: The T cell’s license to kill stem cells in the inflamed intestine. Sci Immunol 4: 2019.

35. Miyoshi H, Ajima R, Luo CT, Yamaguchi TP, and Stappenbeck TS. Wnt5a potentiates TGF-beta signaling to promote colonic crypt regeneration after tissue injury. Science 338: 108–113, 2012.

36. Howe KL, Lorentz RJ, Assa A, Pinnell LJ, Johnson-Henry KC, and Sherman PM. Transforming growth factor-beta1 protects against intestinal epithelial barrier dysfunction caused by hypoxia-reoxygenation. Shock 43: 483–489, 2015.

37. Matsui T, Ichikawa H, Fujita T, Takemura S, Takagi T, Osada-Oka M, and Minamiyama Y. Histidine and arginine modulate intestinal cell restitution via transforming growth factor-beta1. European journal of pharmacology 2019.

38. Kai K, Iwamoto T, Zhang D, Shen L, Takahashi Y, Rao A, Thompson A, Sen S, and Ueno NT. CSF-1/CSF-1R axis is associated with epithelial/mesenchymal hybrid phenotype in epithelial-like inflammatory breast cancer. Sci Rep 8: 9427, 2018.

39. Menke J, Iwata Y, Rabacal WA, Basu R, Yeung YG, Humphreys BD, Wada T, Schwarting A, Stanley ER, and Kelley VR. CSF-1 signals directly to renal tubular epithelial cells to mediate repair in mice. Journal of Clinical Investigation 119: 2330–2342, 2009.

40. Huynh D, Dai XM, Nandi S, Lightowler S, Trivett M, Chan CK, Bertoncello I, Ramsay RG, and Stanley ER. Colony stimulating factor-1 dependence of paneth cell development in the mouse small intestine. Gastroenterology 137: 136–144, 144 e131-133, 2009.

41. Chambers SK. Role of CSF-1 in progression of epithelial ovarian cancer. Future Oncology 5: 1429–1440, 2009.

42. Huynh D, Akçora D, Malaterre J, Chan CK, Dai X-M, Bertoncello I, Stanley ER, and Ramsay RG. CSF-1 Receptor-Dependent Colon Development, Homeostasis and Inflammatory Stress Response. PLoS ONE 8: e56951, 2013.

43. Akcora D, Huynh D, Lightowler S, Germann M, Robine S, de May JR, Pollard JW, Stanley ER, Malaterre J, and Ramsay RG. The CSF-1 receptor fashions the intestinal stem cell niche. Stem Cell Res 10: 203–212, 2013.

