## Supplementary figures and images for "Single-cell transcriptomics predict novel potential regulators of acute epithelial restitution in the ischemia-injured intestine"

### Supplemental Figure 1

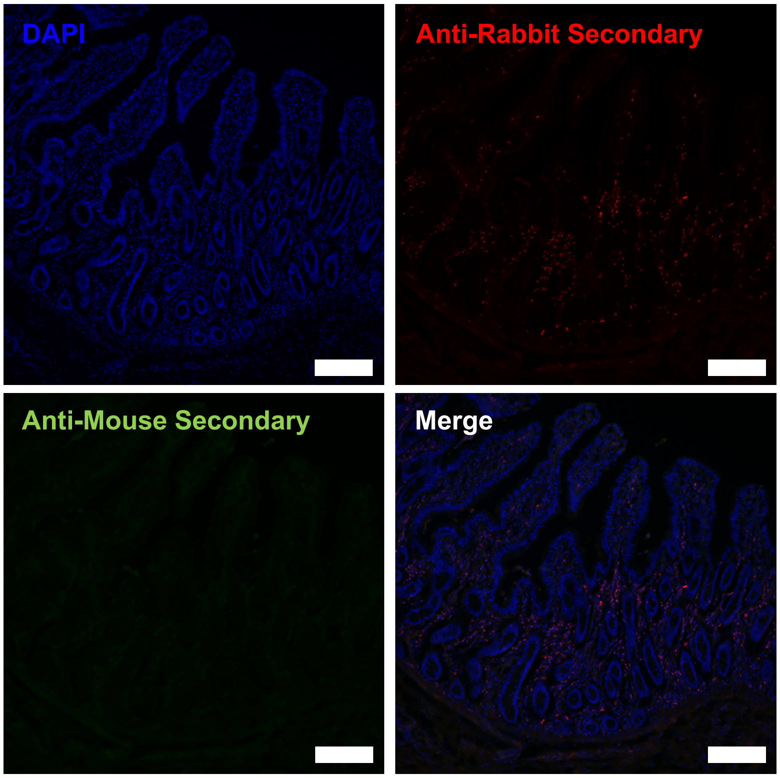
